# EFMouse: a toolbox to model stimulation-induced electric fields in the mouse brain

**DOI:** 10.1101/2024.07.25.605227

**Authors:** Ruben Sanchez-Romero, Sibel Akyuz, Bart Krekelberg

**Affiliations:** Center for Molecular and Behavioral Neuroscience, Rutgers University, Newark, NJ, 07102, USA

**Author notes:** Corresponding author (R. Sanchez-Romero).

**Keywords:** electrical stimulation, electric field modeling, mouse models, high-definition stimulation, transcranial direct current stimulation

## Abstract

Research into the mechanisms underlying neuromodulation by tES using in-vivo animal models is key to overcoming experimental limitations in humans and essential to building a detailed understanding of the in-vivo consequences of tES. Insights from such animal models are needed to develop targeted and effective therapeutic applications of non-invasive brain stimulation in humans. The sheer difference in scale and geometry between animal models and the human brain contributes to the complexity of designing and interpreting animal studies. Here, we introduce EFMouse, a toolbox that extends previous approaches to model intracranial electric fields and generate predictions that can be tested with in-vivo recordings in mice. Novel functionality includes the ability to capture typical surgical approaches in the mouse (e.g., cranial recording windows), the placement of stimulation electrodes anywhere in or on the animal, and novel ways to report field predictions, including some refined measures of focality and direction homogeneity, and quantification based on regions defined in the Allen Mouse Brain Atlas. Although the EFMouse toolbox is generally applicable to planning and designing tES studies in mice, we illustrate its use by posing questions about transcranial direct current stimulation (tDCS) experiments with the goal of targeting the left visual cortex of the mouse. The EFMouse toolbox is publicly available at https://github.com/klabhub/EFMouse.

**Author summary:** Transcranial electrical stimulation offers opportunities for studying brain activity and developing neuromodulation therapies. Even though this technique is used extensively in humans, understanding its neural consequences is still limited. Mouse models offer an opportunity to bridge this gap. However, major differences between human and mouse brains, such as brain size and cortical folding, pose a challenge to the design of appropriate stimulation protocols in mice. To address this, we developed EFMouse, an open-source computational toolbox that predicts intracranial electrical fields in the mouse brain during stimulation. This toolbox allows researchers to design experiments by simulating electrode arrangements and quantifying properties of the predicted electric field in specific brain regions. By doing so, EFMouse can guide the optimization of stimulation techniques to achieve targeted and reproducible effects. We illustrate its use by comparing a series of electrode arrangements, in terms of the strength, focality, and direction of their induced electric field. By making EFMouse publicly available, we hope to advance fundamental neuroscience research and the development of future clinical applications.

**Highlights:**

EFMouse is a novel, open-source, Matlab-based electric field simulator for the mouse brain.
EFMouse quantifies induced field focality and homogeneity in regions of the Allen Mouse Brain Atlas.
Montages with a return on the mouse’s back generate homogeneous fields perpendicular to the cortical surface.
Montages with a small distance between stimulation and return electrodes on the mouse’s head can generate focal, but relatively weak fields.

## Introduction

Understanding intracranial current flow is an essential aspect of harnessing the potential of transcranial current stimulation. In-vivo intracranial measurements provide the most direct assessment of intracranial fields caused by tES. [1–3] provide such measurements of electric field magnitudes in humans. While important, such studies with human volunteers are limited because recording electrodes are placed based on clinical considerations and not to test the properties of induced electric fields. These limitations are partially addressed by translational work in animal models [1,4–8] and human cadavers [9,10], although these come with a different set of caveats [11]. Reviews in [12] and [13] provide a list of considerations to better translate electrical stimulation experiments in rodents and non-human primates to humans, for example, differences in brain geometry that determine the flow of electric current, orientation of targeted neuronal columns and placement of stimulation electrodes, organization of functional networks related to targeted behavior, and the need to adjust parameters, such as the injected current, to achieve field magnitudes that could realistically be achieved in human applications. Aspects such as axonal length [14], higher neuronal densities observed in primates [15] and network effects [16,17], are beyond the scope of the current work, but are also important to consider for the design and interpretation of electrical stimulation experiments in animal models.

Given the technical challenges of intracranial field recordings, electric field models provide a complementary approach to investigating various conditions of interest. For instance, [18] used high-resolution anatomical mesh models to compare electric fields across mouse, monkey, and human brains, and [19] used the same anatomical mouse mesh to simulate a montage with one electrode clipped in each ear. These studies primarily reported results at the whole-brain level, emphasizing general similarities and differences between species.

Here, we build on these approaches by developing a mouse electric field model (EFMouse) specifically for experiments in which the neural consequences of stimulation are measured invasively, for example, with intracranial recording electrodes or two-photon imaging (e.g., [20]). This required several refinements that were not available in other open-source electric field modeling software packages optimized for non-invasive stimulation of human brains (SimNIBS [21] and ROAST [22]). 1) We modeled the skin removal and the craniotomies that are necessary to probe intracranial responses. For example, in EFMouse, users can model a cranial recording window and analyze the predicted electric field in the underlying cortical region. This feature increases the validity of the model predictions by considering surgery-induced tissue inhomogeneities affecting the current flow. 2) We developed an approach that allows the stimulation electrodes to be placed in any tissue or region of the animal’s body. For instance, users can model stimulation electrodes positioned intracranially, on the skin or bone of the mouse head or other commonly used body parts, such as the lumbar region, submandibular area, or neck. 3) We created tools to quantify field properties in arbitrary regions of interest defined by stereotaxic coordinates or in identified regions of the Allen Mouse Brain Atlas. Users can quantify the electric field on a whole-brain basis, compare fields in target (e.g., under the cranial window) and off-target regions of interest, or use the Allen atlas to analyze the field properties across pre-defined functional areas. These novel functionalities bring us closer to a model that informs questions about target and off-target stimulation, enables the comparison of the relative benefits of different montages, and ultimately predicts specific (neural or behavioral) outcomes in an experiment.

This paper first presents the considerations and technical solutions that led to EFMouse (Design and Implementation). Second, we illustrate its use by asking questions about planned studies in mouse visual cortex. Separately, we model noninvasive experiments in which stimulation electrodes are placed on the scalp (without surgical intervention, as might be the case in an experiment investigating only the behavioral consequences of tES) and invasive experiments in which a craniotomy is created to measure neural consequences of stimulation. Researchers can adapt EFMouse to their questions following the MATLAB tutorials in the project repository (https://github.com/klabhub/EFMouse).

### Design and Implementation

The toolbox was implemented using an object-oriented approach in MATLAB (R2023a). The code is documented extensively, includes tutorials matching the simulations presented in this paper, and is available at https://github.com/klabhub/EFMouse. All parameters in EFMouse are configurable, allowing users to adjust them at will. For details of the user interface, we refer to the tutorials and code, but we provide a coarse overview here. A typical workflow consists of five stages: INIT, MESH, EXPORT, GETDP, and ATLAS. In the

INIT stage, EFMouse creates a folder to store (intermediate) results and log files, allowing users to restart a simulation or inspect intermediate results. After defining stimulation electrodes and, optionally, a craniotomy, the MESH stage generates the finite element mesh. EFMouse offers various tools to visualize the mesh, which should be used to confirm the placement of electrodes and craniotomies. The EXPORT stage creates text files for use with the finite element solver GetDP and the GETDP stage uses these files to solve the electrostatic problem. Once GetDP finishes its simulations, the results can be visualized in the mesh, or transverse, sagittal or coronal sections through the brain. Analysis of field strength, focality, and homogeneity can be based on arbitrarily defined regions of interest, or regions of the Allen atlas with the ATLAS stage (for the latter, FSL 6.0.7 onwards is required (https://fsl.fmrib.ox.ac.uk/fsl/)).

### Mouse mesh

We use the mouse mesh published in [18], based on images from the Digimouse project [23]. In short, the mesh corresponds to one 28-gram normal nude male mouse for which post-mortem isotropic 0.1mm voxel images were acquired (CT, PET, and cryosection). The high-resolution mesh provided in [18] consists of approximately 1M nodes and 5.7M tetrahedra and segments the mouse’s whole body into five broad structures with corresponding conductivity values: gray matter (0.275 Siemens per meter, S/m), CSF (1.654 S/m), bone (0.01 S/m), soft tissue (0.465 S/m) and eyeballs (0.5 S/m). We changed the mesh’s orientation to left to right (x-axis), posterior to anterior (y-axis), and inferior to superior (z-axis) to match the conventions of the Paxinos atlas [24].

### Craniotomy

Measuring neural consequences of stimulation at the single neuron level requires an invasive approach and, therefore, a craniotomy. EFMouse creates a circular or rectangular craniotomy by assigning a new material type (i.e., a conductivity) to the mesh elements representing the bone and skin that are surgically removed. The user is free to choose the conductivity that best reflects the surgical approach. In the simulations below we explore the difference between a surgical approach that leaves a dry craniotomy (i.e., skin and bone tetrahedra are replaced with air (2.5e-14 S/m)) and an approach in which the craniotomy fills with a conducting material (e.g., saline or CSF).

### Electrode placement

In tES experiments, electrodes are typically placed on top of the skin. In EFMouse, these electrodes are called *surface* electrodes and modeled as a cylindrical or cuboid-shaped mesh that is locally orthogonal to the mouse’s skin. Because they are constructed by orthogonal extrusion from the skin surface, the resulting 3D shape is not perfectly cylindrical or cuboid (especially for larger electrodes) and is best thought of as a conducting gel that molds to the skin’s surface. In the simulations below, we used circular surface electrodes of 0.6 mm radius and 1 mm thickness, and rectangular electrodes of 15 mm width × 15 mm length × 1 mm thickness. The conductivity of the electrode tetrahedra was set to 0.3 S/m (i.e., electrode gel). Implanted electrodes are often used for chronic recordings in awake animals. In EFMouse, these are called *insert* electrodes and modeled by setting the conductivity of a cylindrical or cuboid region in the mouse mesh to a high value. This is intended to capture an approach where, for instance, a metal screw is inserted into the cranial bone.

### Guiding placement

The mesh’s anatomical resolution does not provide detailed landmarks. To improve the placement of instrumentation (e.g., the craniotomy and the electrodes), we manually coregistered the Digimouse’s volumetric NIfTI files with the visual areas defined in the Allen atlas (see **Fig 5** for a volume section example). EFMouse provides a function to map from volumetric coordinates to mesh coordinates. (Digimouse and Allen atlas NIfTI files are provided in the repository.)

### Electric field modeling

The finite element method (FEM) solution for the electric field problem described by the mouse, craniotomy, electrodes, conductivities, and injected currents, is based on the ROAST implementation but adjusted to the specifics of our mouse model. We solved the Laplace equations with Neumann boundary conditions imposed on the electrodes using the open-access solver GetDP v.3.5.0 [25] (https://getdp.info/). This software computes the voltage (V), electric field components in x, y and z directions (V/m), and electric field magnitude (V/m) at each node in the mesh.

### Relative focality

Extending the definition of focality used in [26], we define a highly focal montage as one in which few areas outside the target area have *eMag* values as high as the ones observed in the target area. Such a montage produces a strong electric field concentrated in the target area. Low focality indicates the opposite: many reference areas have *eMag* values as high as the target area—a montage with more diffuse effects. To capture this, we defined relative focality as: one minus the fraction of nodes in a *reference area* with an electric field magnitude (*eMag*) greater than *P* of the *target area’s* 99.9th percentile of its *eMag* distribution. Our measure of relative focality ranges from zero to one. Values close to zero indicate low relative focality, values close to one indicate high relative focality. In our simulations, we observed that a *P* = 40% enhances sensitivity to diffuse effects; however, users can adjust *P* to suit their research questions.

### Direction homogeneity

In-vitro [27,28] and in-vivo [29] recordings show the importance of the direction of the induced field relative to the orientation of the major axis of the neurons. Based on such results, a field with a small magnitude aligned homogeneously with the neurons could have a bigger influence than a misaligned field with a large magnitude. We define direction homogeneity as the magnitude of the mean of the normalized components across all nodes in a given region.

Direction homogeneity is thus computed as sqrt(mean(*eX_n*)^2^ + mean(*eY_n*)^2^ + mean(*eZ_n*)^2^), where *eX_n* = *eX*/*eMag* is the normalized field component in the x direction for each mesh node, equivalently for *eY_n* and *eZ_n*. Normalized components are helpful in illustrating the direction of the field irrespective of its magnitude. Also, by using normalized components, direction homogeneity ranges from 0 to 1, with larger values indicating more homogeneity and thus electric field components with a more uniform direction. (Consider an extreme example where half of the nodes point to the positive direction and the other half to the negative direction; the resulting field direction homogeneity will be zero.) In the Results section, we report direction homogeneity and mean normalized x, y, and z components.

## Results

We developed a MATLAB-based toolbox (EFMouse) to simulate intracranial fields in the mouse brain induced by current stimulation. The GitHub repository (https://github.com/klabhub/EFMouse) contains tutorials and tools that allow researchers to ask questions geared to their experiments. Here, we illustrate its use by posing questions about transcranial direct current stimulation (tDCS) experiments to target the left visual cortex. We explore the role of electrode montages, craniotomies, and quantification across the whole brain, arbitrarily shaped regions of interest, and regions of the Allen Brain Atlas in terms of average field strengths, homogeneity and focality.

### Standard versus high-definition montages

Our first question was whether a high-definition, center-surround montage [30,31] is advantageous over a traditional two-electrode montage, as has been claimed based on human behavioral data [32,33]. To explore this question in a mouse model, we simulated a standard montage (referred to as 1 x Back) with one circular anode (+0.2 mA) targeting the left visual cortex and a larger rectangular cathode (-0.2 mA) placed on the mouse’s back following [29,34] (**Fig 1A**); and a high-definition montage (referred to as 1 × 4) with one circular anode (+0.2 mA) above visual cortex and four circular cathodes (-0.05 mA each) surrounding the anode (**Fig 1B**).

**Fig 1.**
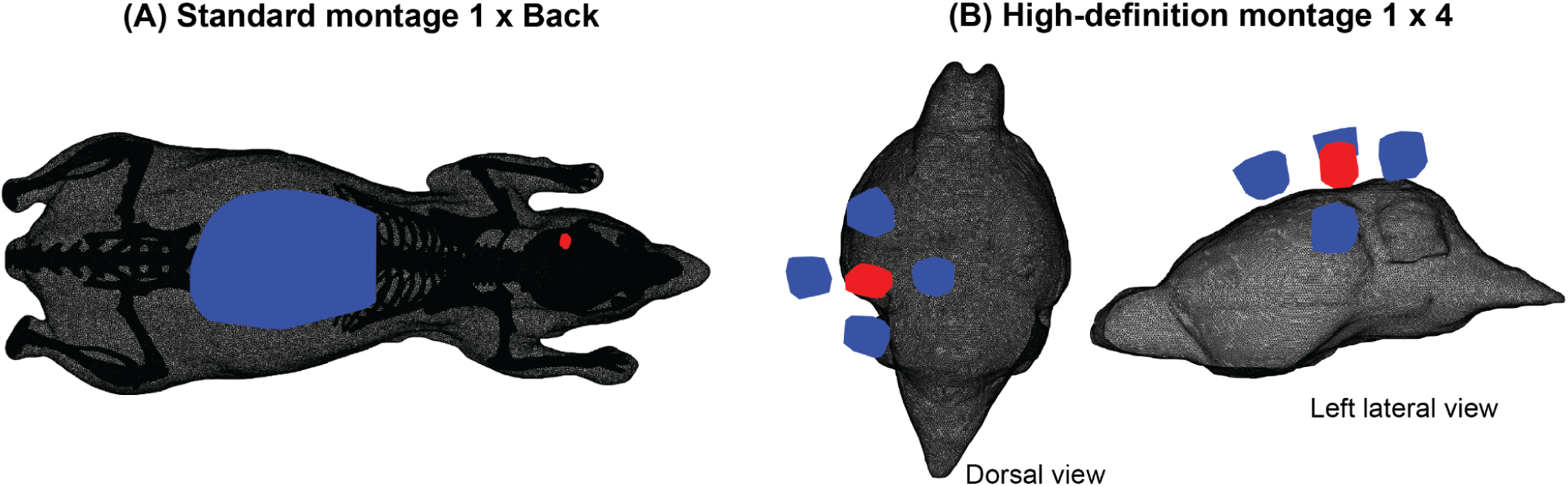
Standard and high-definition electrical stimulation montages. Visualization in mesh space showing mouse tissue (black), anodes (red), and cathodes (blue). (A) Standard montage 1 x Back. The surface electrodes are not perfectly rectangular or cylindrical because they follow the local shape of the skin (Design and Implementation). (B) High-definition montage 1 × 4, zoomed in to the mouse brain.

We analyzed the field at the whole-brain level. Specifically, we quantified fields at the nodes labeled as “gray matter” in the Digimouse/Alekseichuk mesh; this contains olfactory bulbs, cortex, subcortex, cerebellum, and the brainstem. EFMouse can also produce electric field summary statistics for the other mesh tissues/elements, including soft tissue, bone, CSF, eyeballs, craniotomy, and electrodes. The MATLAB tutorials in the GitHub repository provide examples. **Fig 2** visualizes the field across the brain for the 1 x Back montage (top panels) and three orthogonal sections through the mouse brain (bottom panels). Visual inspection of **Fig 2** suggests that this montage targets the left visual cortex as intended: the area surrounding the stimulation electrode has a higher field magnitude than the contralateral right area.

**Fig 2.**
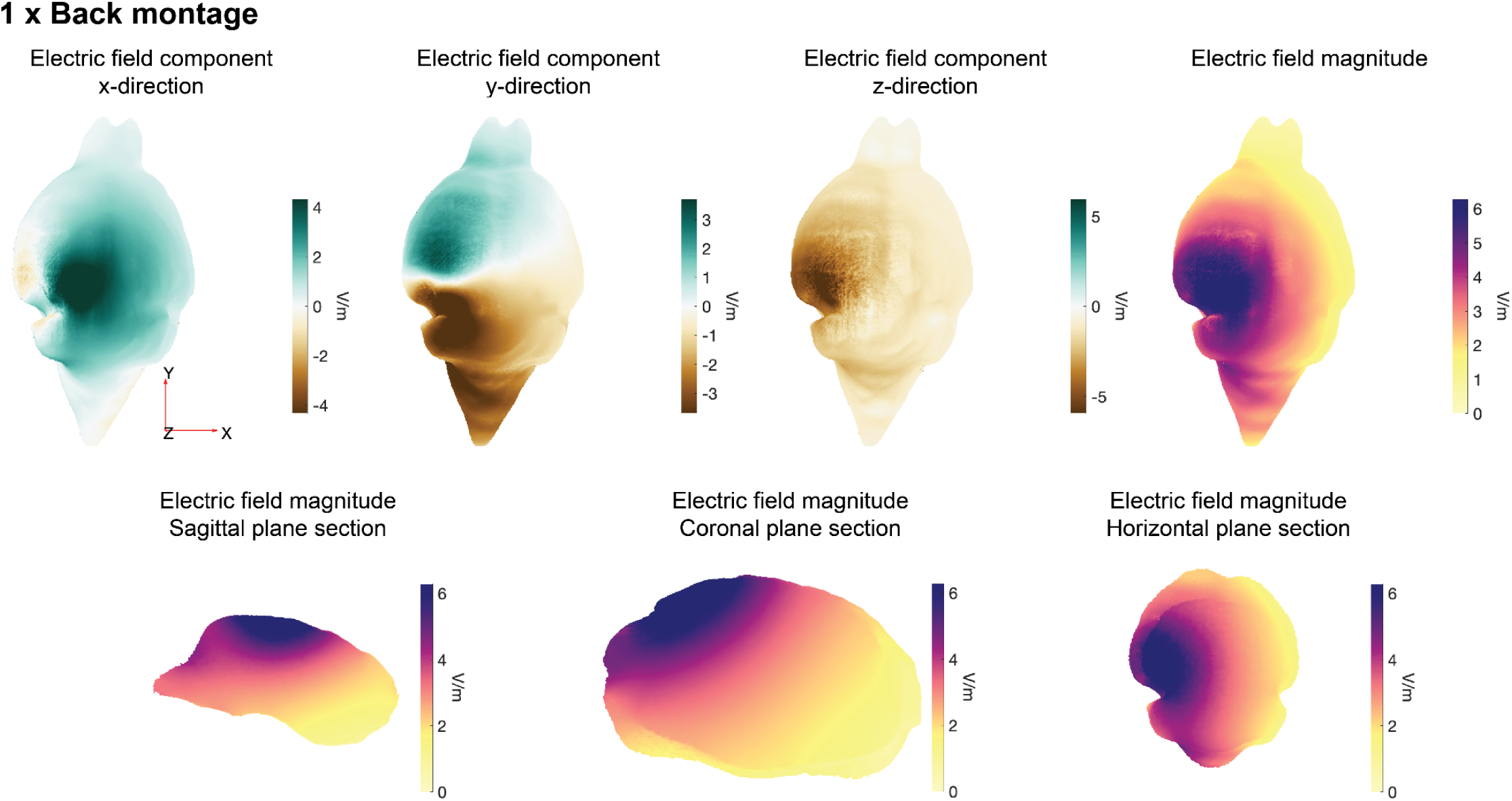
Predicted electric field (V/m) in the whole Digimouse brain. Visualization in mesh space. Results for 1 x Back montage. Field magnitudes are capped at the 98th percentile to remove outliers from this visualization. Top panels show 3D views. Bottom panels show field magnitudes for three orthogonal sections.

**Table 1A** quantifies the whole-brain fields, showing that the 1 x Back montage produces a substantial component in the z direction. This component is of particular interest as it aligns approximately with the orientation of pyramidal neurons in visual cortex and is therefore expected to have the largest neural effects [28,35,36]. The field was also relatively homogeneous. **Table 1B** quantifies the whole-brain fields produced by the high-definition 1 × 4 montage. Notably, these magnitudes are almost five times smaller than the 1 x Back montage, suggesting that the tight arrangement of the high-definition montage shunts most of the current through the skin. In addition, the direction homogeneity of the 1 × 4 montage was also lower.

**Table 1.**
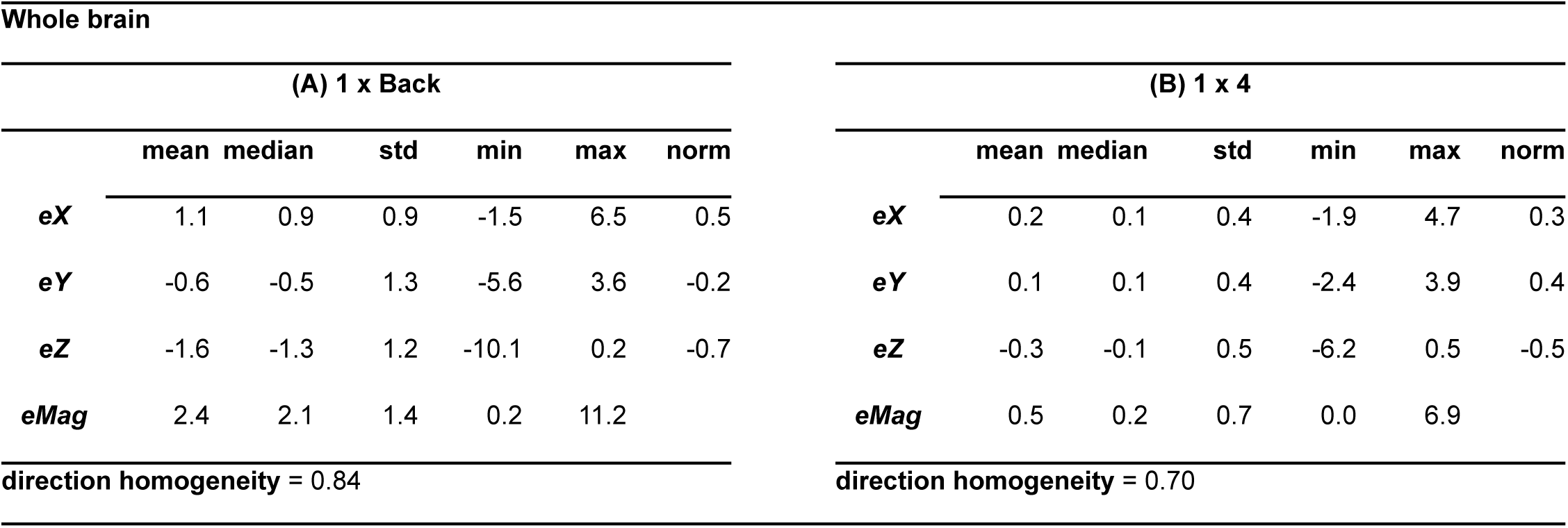
Summary statistics for the predicted electric field (V/m) in the Digimouse brain. Results for the x, y, and z direction field components (*eX*, *eY,* and *eZ*), field magnitude (*eMag*) and direction homogeneity for the (A) 1 x Back, and (B) 1 × 4 montages. The norm column represents the mean normalized field components. (Rounding may show some values as zero.)

Whole brain
| (A) 1 x Back |  |  |  |  |  |  | (B) 1 x 4 |  |  |  |  |  |  |
| --- | --- | --- | --- | --- | --- | --- | --- | --- | --- | --- | --- | --- | --- |
|  | mean | median | std | min | max | norm |  | mean | median | std | min | max | norm |
| **eX** | 1.1 | 0.9 | 0.9 | -1.5 | 6.5 | 0.5 | **eX** | 0.2 | 0.1 | 0.4 | -1.9 | 4.7 | 0.3 |
| **eY** | -0.6 | -0.5 | 1.3 | -5.6 | 3.6 | -0.2 | **eY** | 0.1 | 0.1 | 0.4 | -2.4 | 3.9 | 0.4 |
| **eZ** | -1.6 | -1.3 | 1.2 | -10.1 | 0.2 | -0.7 | **eZ** | -0.3 | -0.1 | 0.5 | -6.2 | 0.5 | -0.5 |
| **eMag** | 2.4 | 2.1 | 1.4 | 0.2 | 11.2 |  | **eMag** | 0.5 | 0.2 | 0.7 | 0.0 | 6.9 |  |
| direction homogeneity = 0.84 |  |  |  |  |  |  | direction homogeneity = 0.70 |  |  |  |  |  |  |

### Craniotomies

Above, we simulated an experiment in which stimulation was applied to an animal with a fully intact skull. This could, for instance, help to interpret the results of an experiment in which the effects of tDCS are assessed behaviorally (e.g., in this montage, one might predict changes in the animal’s visual sensitivity in the right visual field). However, a key goal of animal experiments is to measure the neural consequences of tES directly, at the single neuron level. Such experiments require physical access to the brain and hence a craniotomy. In this section, we use EFMouse to explore how craniotomies affect the fields in the brain.

We modeled a circular craniotomy placed above the left visual cortex by replacing the corresponding skin and bone tissue with poorly conducting material (2.5e-14 S/m). This was intended to capture a common two-photon imaging setup in which the craniotomy is electrically insulated from the skin. We explored two montages with this craniotomy. In the first, called 4 x Back, we positioned four anodes (+0.05 mA each) surrounding the craniotomy (**Fig 3A**), and a rectangular cathode on the mouse’s back (-0.2 mA). In the second, called 1 × 1 (**Fig 3B**), we placed one anode anterior to the craniotomy (+0.2 mA) and one cathode posterior to the craniotomy (-0.2 mA). **Table 2** shows that these montages produce similar field magnitudes, but qualitatively different orientations. Notably, the 4 x Back montage (**Table 2A**) has a substantial dorsal-to-ventral component (negative component in the z direction); while the 1 × 1 montage has a substantial anterior-to-posterior component (negative component in the y direction).

**Fig 3.**
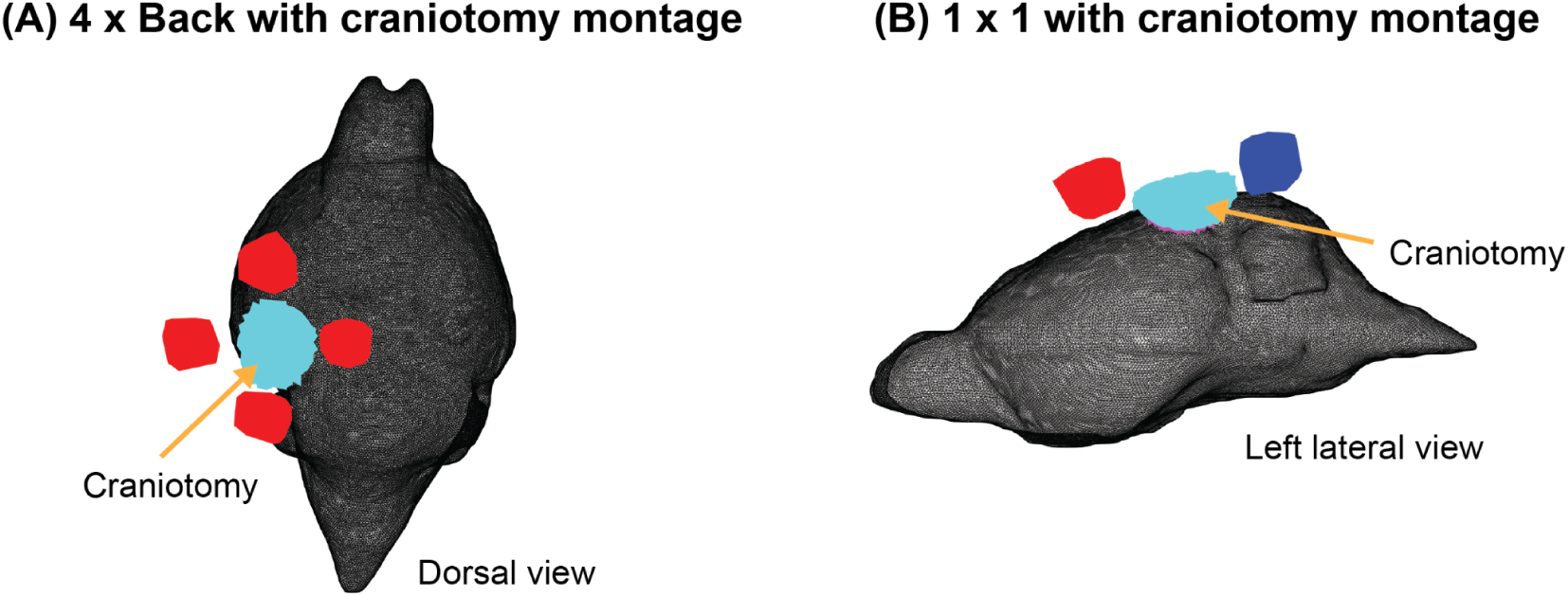
Electrical stimulation montages with craniotomy. Visualization in mesh space showing mouse tissue (black), anodes (red), cathodes (blue), removed skin (cyan), and underlying removed bone (magenta) from the craniotomy. (A) 4 x Back montage, zoomed in to the mouse brain. The cathode in the mouse’s back is not shown. (B) 1 × 1 montage. The surface electrodes are not perfectly rectangular or cylindrical because they follow the local shape of the skin (Design and Implementation).

**Table 2.** Summary statistics for the predicted electric field (V/m) in the Digimouse brain for montages with craniotomy. Results for the x, y, and z direction field components (*eX*, *eY,* and *eZ*), field magnitude (*eMag*) and direction homogeneity for the (A) 4 x Back with craniotomy, and (B) 1 × 1 with craniotomy montages. The norm column represents the mean normalized field components.

Whole brain
| (A) 4 x Back with craniotomy |  |  |  |  |  |  | (B) 1 x 1 with craniotomy |  |  |  |  |  |  |
| --- | --- | --- | --- | --- | --- | --- | --- | --- | --- | --- | --- | --- | --- |
|  | mean | median | std | min | max | norm |  | mean | median | std | min | max | norm |
| $eX$ | 0.9 | 0.8 | 0.5 | -1.7 | 3.0 | 0.5 | $eX$ | 0.7 | 0.7 | 1.4 | -7.8 | 7.1 | 0.2 |
| $eY$ | -0.8 | -0.7 | 1.0 | -3.4 | 1.8 | -0.3 | $eY$ | -2.1 | -1.8 | 2.0 | -13.7 | 2.6 | -0.7 |
| $eZ$ | -1.3 | -1.2 | 0.7 | -4.7 | 0.3 | -0.6 | $eZ$ | -0.6 | -0.4 | 1.8 | -11.5 | 7.6 | -0.2 |
| $eMag$ | 2.0 | 2.0 | 0.9 | 0.2 | 5.3 | | $eMag$ | 3.1 | 2.4 | 2.2 | 0.2 | 15.1 | |
| direction homogeneity = 0.86 |  |  |  |  |  |  | direction homogeneity = 0.74 |  |  |  |  |  |  |

Interestingly, comparing the results for the 4 x Back montage with craniotomy (**Table 2A**) with the 1 x Back montage without a craniotomy (**Table 1A**) shows that they were practically equivalent, as were the results for homogeneity. This suggests that at the whole-brain level, the simulated craniotomy has relatively little influence on the intracranial fields.

The craniotomy we modeled assumed a very low conductivity path (air) from the electrodes on the skin to the brain. In some experiments, however, a high conductivity path could be formed by saline or CSF filling the craniotomy and bridging the air gap between brain and skin. We investigated the potential impact of such a shunt in the 1 × 1 montage by replacing the bone and skin tissue removed for the craniotomy with a highly conducting material (1.654 S/m). The main effect was an overall reduction in the field magnitude by about 30% (compared to the dry craniotomy simulated above), but neither the dominant orientation, nor the homogeneity changed substantially (Results not shown, but available in the GitHub repository tutorials).

### Regions of Interest analysis

The analyses reported above provide summary information on fields in the brain. In an experiment that aims to assess neural consequences of tES, however, the fields produced in a specific region of interest—the region where neural activity is recorded—are of primary interest. To address this, EFMouse provides functionality to quantify electric fields in a box-shaped region of interest (ROI).

Here we defined one ROI centered in the craniotomy in the left hemisphere (with dimensions width = 2 mm, length = 2 mm, depth = 1 mm) (**Fig 4A**), and for comparison, an analogous ROI in the right hemisphere with matching dimensions (**Fig 4B**). Results in **Table 3A-B** show that for the ROI under the craniotomy, (i) the 1 × 1 montage produced a stronger mean magnitude; (ii) the 4 x Back field generated a relatively stronger dorsal-to-ventral field (mean *eZ* = -2.2 V/m), while the field produced by the 1 × 1 montage was primarily in the anterior-to-posterior y direction (mean *eY* = -6.4 V/m). Results in **Table 3C-D** for the comparison ROI in the right visual cortex show substantially smaller (but still non-zero) fields.

**Fig 4.**
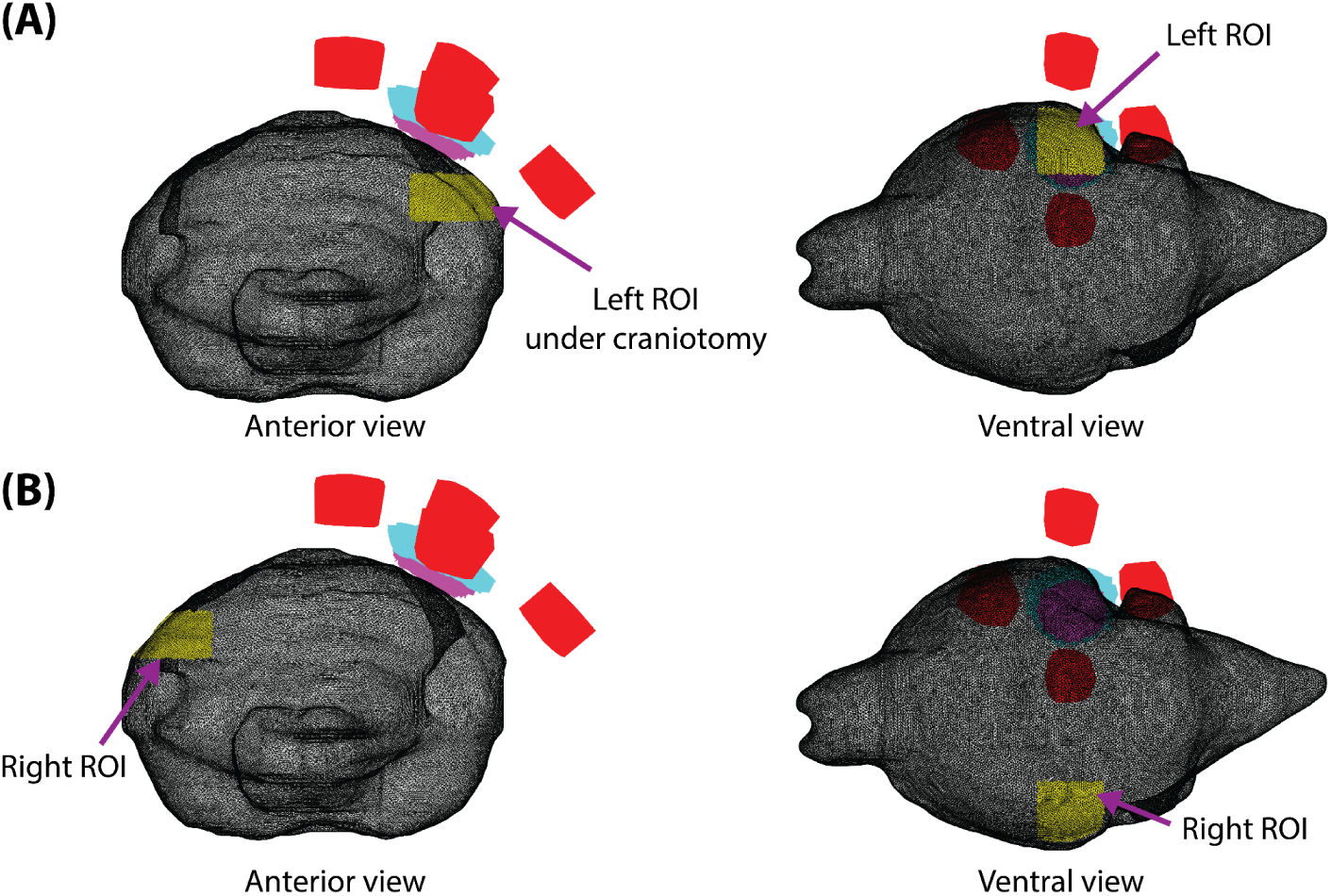
Regions of interest (ROIs). Visualization in mesh space. (A) A target ROI is located in the left visual cortex under the craniotomy. (B) The comparison ROI with matching dimensions is in the contralateral right visual cortex. Electrodes and craniotomy for the 4 x Back montage only. ROIs are depicted in yellow, anodes in red, removed skin in cyan, removed bone in magenta. The cathode is not shown. EFMouse can provide summary statistics on any ROI, enabling a comparison of on-target and off-target fields.

**Table 3.** Summary statistics for the predicted electric field (V/m) for two box-shaped regions of interest (ROIs) in the mouse visual cortex. Results for (A-B) ROI under craniotomy in the left visual cortex and (C-D) analogous ROI in the contralateral right visual cortex. Results for the x, y, and z direction field components (*eX*, *eY,* and *eZ*), field magnitude (*eMag*), direction homogeneity and relative focality, for the 4 x Back (A,C) and 1 × 1 (B,D) montages. The norm column represents the mean normalized field components.

| ROI under craniotomy in left visual cortex |  |  |  |  |  |  |  |  |  |  |  |  |  |
| --- | --- | --- | --- | --- | --- | --- | --- | --- | --- | --- | --- | --- | --- |
| (A) 4 x Back with craniotomy |  |  |  |  |  |  | (B) 1 x 1 with craniotomy |  |  |  |  |  |  |
|  | mean | median | std | min | max | norm |  | mean | median | std | min | max | norm |
| *eX* | 1.5 | 1.5 | 0.3 | 0.2 | 2.1 | 0.5 | *eX* | 2.1 | 2.0 | 1.2 | -1.1 | 6.5 | 0.3 |
| *eY* | -1.5 | -1.5 | 0.3 | -2.5 | -0.5 | -0.5 | *eY* | -6.4 | -6.4 | 1.5 | -11.3 | -2.6 | -0.9 |
| *eZ* | -2.2 | -2.2 | 0.4 | -3.3 | -0.9 | -0.7 | *eZ* | -1.8 | -1.5 | 1.6 | -6.6 | 2.7 | -0.2 |
| *eMag* | 3.1 | 3.1 | 0.3 | 2.1 | 3.9 |  | *eMag* | 7.3 | 7.2 | 1.5 | 4.2 | 11.7 |  |
| direction homogeneity = 0.98 |  |  |  |  |  |  | direction homogeneity = 0.96 |  |  |  |  |  |  |

| (C) 4 x Back with craniotomy |  |  |  |  |  |  | (D) 1 x 1 with craniotomy |  |  |  |  |  |  |
| --- | --- | --- | --- | --- | --- | --- | --- | --- | --- | --- | --- | --- | --- |
|  | mean | median | std | min | max | norm |  | mean | median | std | min | max | norm |
| eX | 1.1 | 1.1 | 0.2 | 0.5 | 1.6 | 0.6 | eX | 0.4 | 0.4 | 0.4 | -0.9 | 1.3 | 0.2 |
| eY | -0.9 | -0.8 | 0.2 | -1.5 | -0.5 | -0.5 | eY | -2.0 | -2.0 | 0.3 | -3.1 | -1.3 | -1.0 |
| eZ | -1.1 | -1.1 | 0.1 | -1.4 | -0.8 | -0.6 | eZ | -0.2 | -0.2 | 0.1 | -0.5 | 0.2 | -0.1 |
| eMag | 1.8 | 1.8 | 0.2 | 1.3 | 2.3 |  | eMag | 2.1 | 2.1 | 0.3 | 1.4 | 3.1 |  |
| direction homogeneity = 0.99 |  |  |  |  |  |  | direction homogeneity = 0.98 |  |  |  |  |  |  |
| relative focality = 0.09 |  |  |  |  |  |  | relative focality = 0.08 |  |  |  |  |  |  |

The simulations illustrate that tES can produce substantial but widely spread fields in the mouse brain. To quantify the spread of the field, we define a measure called relative focality that compares the field in a target area to a reference area (Design and Implementation). For example, with the ROI under the craniotomy as the *target area* and the rest of the brain as the *reference area* (160,508 nodes), the 4 x Back montage relative focality was 0.35 (**Table 3A**). This implies that 65% of the brain outside the target area received a field similar to that produced inside the target area. The relative focality for the 1 × 1 montage was 0.84 (**Table 3B**), showing that with this montage, only 16% of the brain received a field similar to that produced inside the target area. The analogous computation for the comparison ROI in the right visual cortex results in a relative focality of 0.09 (4 x Back) and 0.08 (1 × 1) (**Table 3C-D**). This shows that almost all the brain (> 91%) received a field as large as the field in the right visual cortex ROI.

### Allen atlas analysis

The ROI-based analysis above disregards anatomical and functional boundaries. Experimental questions, however, are more often formulated with respect to functionally meaningful regions of interest. EFMouse can quantify electric field properties using defined regions in the Allen Mouse Brain Atlas to enable such analyses. To illustrate this, we performed the equivalent analysis of **Table 3**, now using the regions of interest defined by the “visual areas” of the Allen atlas (**Fig 5**). Results in **Table 4A-B** for the left visual cortex are consistent with (and complement) the ROI results from **Table 3A-B**: (i) the 1 × 1 montage produces a stronger field than the 4 x Back; (ii) the 4 x Back montage produces a relatively stronger field in the dorsal-to-ventral negative-z direction, while the 1 × 1 montage produces a substantial field oriented along the anterior-to-posterior negative-y direction. In line with results from **Table 3C-D**, **Table 4C-D** shows that both montages produced weaker fields for the visual areas in the right hemisphere.

**Table 4.**
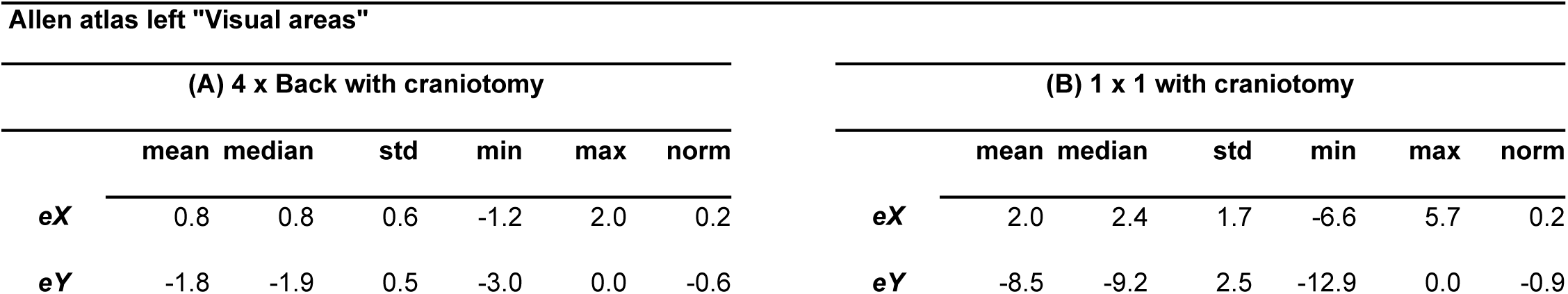
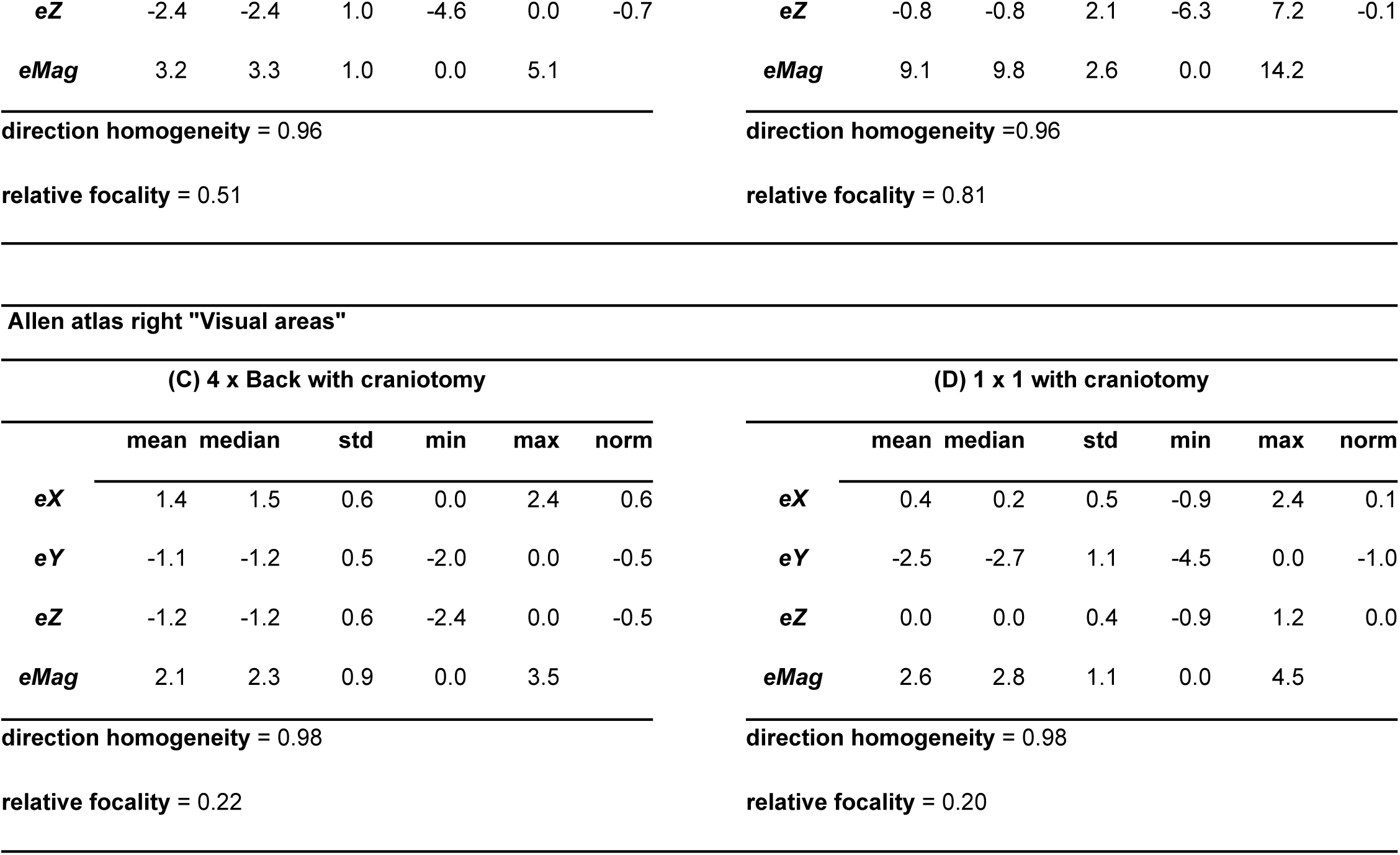
Summary statistics for the predicted electric field (V/m) for the Allen Mouse Brain Atlas visual cortex. Results for (A-B) left visual cortex in the Allen atlas and (C-D) right visual cortex in the Allen atlas. Results for the x, y, and z direction field components (*eX*, *eY,* and *eZ*), field magnitude (*eMag*), direction homogeneity and relative focality, for the 4 x Back (A,C) and 1 × 1 (B,D) montages. The norm column represents the mean normalized field components. (Rounding may show some values as zero.)

**Fig 5.**
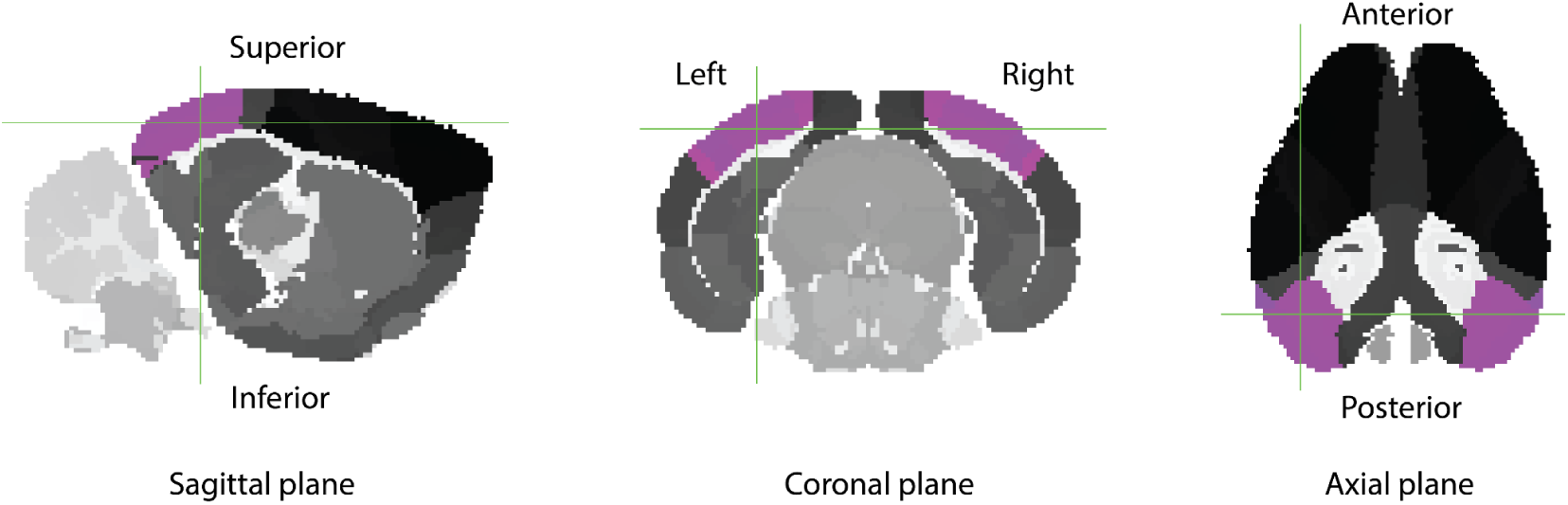
Allen Mouse Brain Atlas visual cortex. The areas labeled “Visual areas” in the Allen atlas are depicted here in magenta. The predicted electric field is analyzed separately for the left and right visual cortex (Table 4). Allen atlas registered to Digimouse volumetric space. Representative section at voxel (28,85,65) in the Allen atlas NIfTI file provided in the EFMouse repository. Image generated with FSLeyes.

Analogous to the box-shaped ROIs, we computed relative focality using the left visual areas of the Allen atlas as the *target area* and the “isocortex” of the Allen atlas (∼92k voxels; excluding the left visual areas) as the *reference area*. Similarly, for the right visual areas. Consistent with the ROI results in **Table 3**, for the left visual cortex, the 4 x Back montage relative focality was 0.51; for the 1 × 1 montage it was 0.81 (**Table 4A-B**). For the right visual cortex, the 4 x Back montage relative focality was 0.22; for the 1 × 1 montage, it was 0.20 (**Table 4C-D**).

## Availability and Future Directions

We developed a toolbox to simulate and analyze electric fields in the mouse brain produced by current stimulation. EFMouse builds on previous work modeling current flow in the brain. We borrowed the mouse anatomical model from [18] and the Digimouse project [23], and the general workflow in the MATLAB implementation from ROAST [22]. We added novel functionality to capture typical surgical approaches in the mouse (e.g., skin removal and cranial recording windows) and the placement of stimulation electrodes anywhere in or on the animal. These enhancements improve the model’s validity for experiments that include intracranial recording (e.g., [37,38]) and bone, epidural, or intracranial stimulation (e.g., [20,39]). In addition, our software adds novel ways to report field predictions, including some refined measures of focality and direction homogeneity, and quantification based on regions defined in the Allen Mouse Brain Atlas. While the model has general applicability, we illustrate its use here by modeling tDCS in mouse visual cortex. We discuss the outcomes and limitations of these analyses and some of the limitations and future directions for the EFMouse model.

Our specific simulations support several conclusions. First, a high-definition, center-surround montage produces considerably smaller and less homogeneous fields than a standard 1 x Back montage. This is likely due to the small distance between the anode and cathodes, which allows the current to be shunted through the skin instead of the brain. Because high-definition montages are intended to generate more localized fields, we assessed relative focality at the ROI and Allen atlas level, and observed that the 1 × 4 montage has higher relative focality compared to the 1 x Back. (Results not shown but available in the Github repository tutorials.) Second, at the whole-brain level, we observed that craniotomies insulated from the skin have only modest effects on the field strengths. However, if the craniotomy forms a conductive bridge between brain and skin, the resulting shunt reduces whole-brain intracranial field strengths. Further complications arise when focusing on specific ROIs, where the craniotomy can increase or decrease the field magnitude, depending on the electrode arrangement. Third, although fields were strongest near the stimulation electrodes, the field in the opposite hemisphere was not negligible. Fourth, montages with a return electrode on the back produced a field in the z direction (coarsely aligned with the principal axis of most neurons in the cortical surface). In contrast, a montage with one return electrode on the head (1 × 1 with craniotomy) generated a field that was predominantly perpendicular to this axis. Because in-vitro and in-vivo recordings show that neural outcomes depend on the alignment of the field with the stimulated neurons [27–29,36], quantifying this alignment could help to interpret tES outcomes. These results point to the more general issue that the induced intracranial fields have complex properties that are not completely represented in brain field magnitude maps (**Fig 2**) or whole-brain level analysis. Interpreting neural or behavioral changes during tES could benefit from a more detailed quantification of the fields. The approaches in EFMouse, including field components and directional homogeneity, comparing on-target and off-target fields with focality, and using coordinate-based ROIs or regions in the Allen Brain Atlas, are a step in that direction.

Relative focality compares field strengths inside a target region with field strengths outside that region. This requires defining a single value that captures the field strength inside the target region (*P*; Design and Implementation). With a high value (e.g., *P* = 100% of the peak inside the target region), relative focality becomes relatively insensitive as it measures only whether nodes outside the target region have fields larger than the peak of the target region. This is unlikely to happen if the target region is near the stimulating electrodes, and relative focality will be 1. With a low *P* value, relative focality also becomes insensitive as all nodes in the model will receive at least some non-zero field, and relative focality becomes 0. In the example simulations, we use *P* = 40%, which provided the sensitivity to detect the differences in focality between the montages we used. However, larger *P* values could be helpful. For instance, if one has reason to assume that field strengths below 90% of the peak in the target region are ineffective, then relative focality using *P* = 90% would identify the off-target regions where stimulation effects should be expected.

One limitation of EFMouse is that the mesh is based on a single nude male mouse from the Digimouse project [23]. This strain differs from the more commonly used C57BL/6 strain, which is also the basis for the Allen atlas. Consequently, the coregistration of the Allen atlas to the Digimouse space will likely be imprecise at the level of the finest functional areas (e.g., cortical layers). If a high-resolution mesh for a C57BL/6 mouse becomes available, it could be incorporated into EFMouse, improving the neuroanatomical interpretation of the predicted electric fields. Similarly, with a suitable mesh, the toolbox could be expanded to other species such as rats, to complement active research combining stimulation and intracranial recordings [4–7,40] and to compare results and methods with previously reported electric field modelling approaches in this species [41,42].

The current version of EFMouse computes the orientation of the electric field in stereotaxic coordinates. For many cortical areas, this can be interpreted as radial or tangential orientations (relative to the cortical surface). However, this is more complex in cortical regions with strong curvature or in subcortical areas (which have not been segmented in the Digimouse mesh). Future work could address this by developing a more detailed mesh of the whole mouse brain.

Finite-element electric field models compute the electric fields based on the well-established principles of electrostatics. However, there are two key factors that produce uncertainty in the model output. First, the conductivities of specific tissue types are not known with great accuracy. Second, the assignment of tissue types to the elements in the model is based on an imperfect segmentation process. For these reasons, one should not expect the predicted fields to match field measurements quantitatively. Studies performing detailed validation of FEM models in humans [3] confirms this; while the large-scale pattern of field magnitudes is well predicted, the specific field magnitudes are not. For these reasons, the absolute numeric values of field strengths generated by EFMouse should be taken with a grain of salt. The value of models such as EFMouse lies more in their qualitative, or relative predictions—for instance, predicting that one montage should result in larger, better-aligned, or more focal fields than another. Such predictions can be tested with measurements of intracranial fields and—with additional assumptions on the modulation of neural activity by applied fields—with measurements of neural and even behavioral changes. Such experiments partially validate the model (and may inform model refinements), but our perspective is that generating testable hypotheses is beneficial in its own right and will ultimately lead to more targeted stimulation protocols and effective interventions in humans.

## Data/code availability statement

All code and data to reproduce our analyses are available at the EFMouse toolbox repository https://github.com/klabhub/EFMouse. EFMouse was implemented in Matlab2023a. For the Allen atlas analysis, FSL 6.0.7 onwards is required (https://fsl.fmrib.ox.ac.uk/fsl/). EFMouse toolbox repository DOI https://doi.org/10.5281/zenodo.14777232 points to the latest version release.

## Author statement

**Ruben Sanchez-Romero**: Conceptualization, Methodology, Software, Formal Analysis, Writing-original draft, Writing-review & editing, Visualization.

**Sibel Akyuz**: Conceptualization, Methodology, Writing-review & editing.

**Bart Krekelberg**: Conceptualization, Methodology, Software, Writing-review & editing, Supervision, Resources, Funding acquisition.

